# Nuclear soluble cGAS senses DNA virus infection

**DOI:** 10.1101/2021.08.27.457948

**Authors:** Yakun Wu, Kun Song, Wenzhuo Hao, Lingyan Wang, Shitao Li

## Abstract

The cytosolic DNA sensor cGAS detects foreign DNA from pathogens or self-DNA from cellular damage and instigates type I interferon (IFN) expression. Recent studies find that cGAS also localizes in the nucleus and binds the chromatin. Despite how cGAS is inhibited in the nucleus is well elucidated, whether nuclear cGAS participates in DNA sensing is not clear. Here, we report that herpes simplex virus 1 (HSV-1) infection caused the release of cGAS from the chromatin into the nuclear soluble fraction. Like its cytosolic counterpart, the leaked nuclear soluble cGAS could sense viral DNA, produce cGAMP, and induce mRNA expression of type I IFN and interferon-stimulated genes. Furthermore, the nuclear cGAS limited HSV-1 infection. Taken together, our study demonstrates that HSV-1 infection releases cGAS from the chromatin tethering and, in turn, the nuclear soluble cGAS activates type I IFN production.

## INTRODUCTION

Cytosolic DNA from infectious microbe triggers innate host defense by activating type I IFN expression (Barber, 2015; Chen et al., 2016; Hornung et al., 2014; Zevini et al., 2017). Cytosolic DNA is recognized by the recently identified DNA sensor, cyclic GMP-AMP synthase (cGAS, also known as MB21D1) (Sun et al., 2013). After binding to DNA, cGAS produces cyclic GMP-AMP (cGAMP) (Sun et al., 2013). cGAMP is a second messenger that binds to the endoplasmic reticulum membrane protein, stimulator of interferon genes (STING, also known as TMEM173, MPYS, MITA, and ERIS), leading to STING dimerization (Ishikawa and Barber, 2008; Sun et al., 2009; Zhang et al., 2013; Zhong et al., 2008). Subsequently, STING recruits TANK-binding kinase 1 (TBK1) to the endoplasmic reticulum and activates TBK1. Activated TBK1 phosphorylates interferon regulatory factors (IRFs), which triggers the dimerization and nuclear translocation of IRFs. In the nucleus, IRFs form active transcriptional complexes and activate type I IFN gene expression (Fitzgerald et al., 2003; Hemmi et al., 2004; McWhirter et al., 2004; Sharma et al., 2003; Tanaka and Chen, 2012).

Excessive host DNA activates the cGAS signaling pathway, leading to aberrant IFN activation and autoimmune diseases, such as Aicardi-Goutieres syndrome (AGS) (Pokatayev et al., 2016). Therefore, cells must render cGAS inert to host genomic DNA, and paradoxically, at the same time, cells need to keep cGAS agile to foreign DNA. The old paradigm is that host DNA is normally restricted to cellular compartments, such as the nucleus and the mitochondria. As cGAS was first thought to be a solely cytosolic protein, the physical barrier blocks the access of cGAS to host DNA (Cai et al., 2014; Chen et al., 2016). However, recent studies further found other subcellular localizations of cGAS, including predominantly in the nucleus (Volkman, 2019), on the plasma membrane (Barnett et al., 2019), mitosis-associated nuclear localization (Gentili et al., 2019; Yang et al., 2017), or phosphorylation-mediated cytosolic retention (Liu et al., 2018). Nonetheless, the nuclear localization of cGAS has been validated by many independent laboratories (Gentili et al., 2019; Mackenzie et al., 2017; Volkman, 2019), which is of particular interest as the nucleus is a DNA-rich environment. Nuclear cGAS is bound to the chromatin in the nucleus by binding to the H2A-H2B dimer of the nucleosome, which immobilizes cGAS on the chromatin; thus, cGAS cannot access the nearby DNA to form an active dimer (Boyer et al., 2020; Cao et al., 2020; Kujirai et al., 2020; Michalski et al., 2020; Pathare et al., 2020; Zhao et al., 2020). In addition, the phosphorylation of the N-terminus of cGAS and the S291 site at the C-terminus further prevents cGAS from activation during mitosis (Li et al., 2021; Zhong et al., 2020). Although the mechanism of nuclear cGAS inhibition is well elucidated, the role of nuclear cGAS in DNA sensing is unknown.

Recent studies showed that the murine R222 (R236 in human cGAS) and murine R241 (R255 in human) sites are the critical sites for the binding to the H2A-H2B dimer (Boyer et al., 2020; Kujirai et al., 2020; Michalski et al., 2020; Pathare et al., 2020; Volkman, 2019; Zhao et al., 2020). The mutation of these two arginines to glutamic acids leads to disruption of the binding to the chromatin and the release of cGAS to nuclear soluble fraction, resulting in cGAS activation and cGAMP production (Volkman, 2019). However, whether and how chromatin-bound cGAS is released into nuclear soluble fraction under physiological or pathological conditions is not clear.

Here, we found that endogenous cGAS tethered with the chromatin in multiple cell lines and HSV-1 infection caused the release of cGAS from the chromatin into the nuclear soluble fraction. The nuclear soluble cGAS bound viral DNA and produced cGAMP. Furthermore, cells exclusively expressing nuclear cGAS responded to HSV-1 infection and activated type I IFN expression. Collectively, our study suggests that HSV-1 infection leads to cGAS release from the chromatin tethering; in turn, the nuclear soluble cGAS senses viral DNA and activates type I IFN to suppress viral infection. Our study uncovers the role of nuclear cGAS in host defense to DNA virus infection.

## RESULTS

### Endogenous cGAS localizes in the cytoplasm and the nucleus

To determine the size of endogenous cGAS protein complex, we performed a sucrose gradient ultracentrifugation for cell lysates from RAW 264.7 macrophages. Unexpectedly, the majority of cGAS proteins were distributed in the high molecular weight fractions with a high sedimentation rate (Fig. 1A). Interestingly, histone H3 co-fractionated with cGAS in most fractions with high molecular weight (Fig. 1A), suggesting that cGAS might associate with the chromatin (Volkman, 2019). To further determine the subcellular localization of endogenous cGAS, we performed subcellular fractionations in multiple cell lines. We fractionated the cell lysates into five fractions: cytosol, membrane, nuclear soluble, chromatin-bound, and cytoskeletal. The nuclear soluble fraction is extracted in a low salt concentration and does not contain histones and nucleosomes, whereas the chromatin-bound fraction comprises nucleosomes with a high salt concentration extract condition. We first fractionated H1299 cell lysates. cGAS was found in the cytosol and the nucleus of H1299 cells (Fig. 1B). Consistent with a previous report (Volkman, 2019), cGAS localized in the chromatin-bound nuclear fraction but not the nuclear soluble fraction (Fig. 1B). Similar results were observed in THP-1 cells (Fig. S1A). We also examined whether DNA stimulation altered cGAS distribution in each fraction. However, the amount of cGAS in each fraction after stimulation was comparable to the fraction without DNA stimulation (Figs. 1B and S1A).

**Figure. 1.**
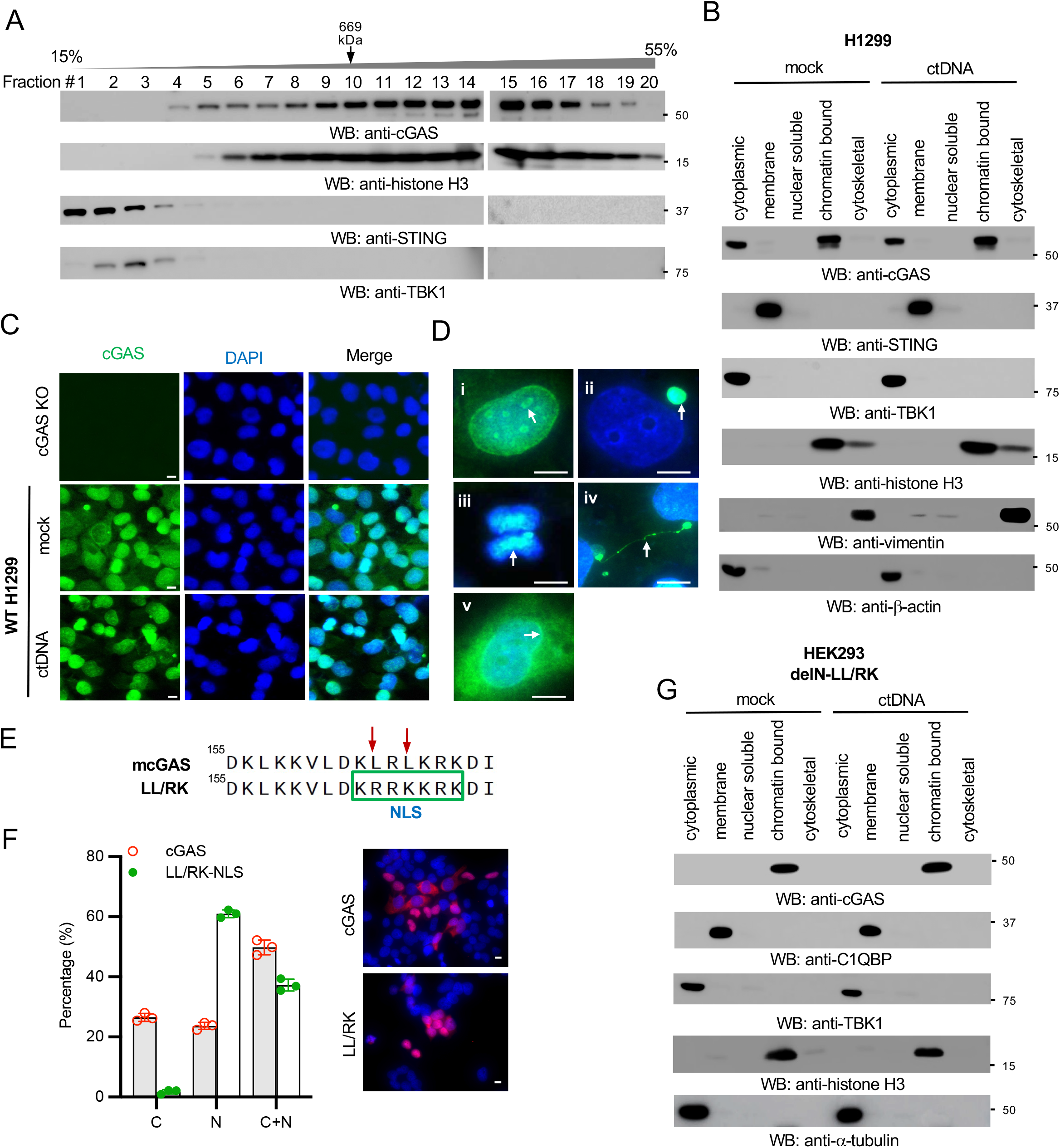
The N-terminal domain and NES regulate cGAS nuclear localization. (**A**) The cell lysates of RAW 264.7 macrophages were separated by 15–55% sucrose density centrifugation. Fractions were blotted as indicated. The fraction of thyroglobulin (660 kDa), a protein standard, was indicated. (**B**) H1299 cells were stimulated with or without 1 μg/ml calf thymus DNA (ctDNA) for 4□h. Then, the cell lysates were fractionated into five fractions: cytoplasmic, membrane, nuclear soluble, chromatin-bound, and cytoskeletal. The fractions were blotted as indicated. STING: membrane marker; TBK1 and β-actin: cytosolic marker; H3: nuclear marker; vimentin: cytoskeletal marker. (**C**) Wild type (WT) and cGAS knockout (KO) H1299 cells were either mock stimulated or transfected with ctDNA. After 4 h, cells were fixed and stained as indicated. cGAS: green; DAPI, blue. Bar = 10 μm. (**D**) Representative cGAS localization in unstimulated H1299 WT cells in (**C**). (**i**) nucleoli; (**ii**) micronucleus; (**iii**) chromosome; (**iv**) chromatin bridge; (**v**) perinuclear region. Arrows indicate each distinct localization in (**i**) to (**v**). cGAS: green; DAPI, blue. Bar = 10 μm. (**E**) Schematic of the LL/RK mutation in the nuclear export signal of cGAS. Red arrows indicate the mutated sites, and the green frame indicates the introduction of a nuclear import signal (NLS) caused by the mutations. (**F**) Immunofluorescence assays of HEK293 cells stably expressing FLAG-tagged mouse cGAS (mcGAS) or the indicated LL/RK mutant. FLAG: red; DAPI, blue. Bar = 10 μm. The summary of the subcellular localization of cGAS and LL/RK mutant was shown in the left panel. C: cytosolic; N: nuclear; C+N: cytosolic and nuclear. Data represent means ± s.d. of three independent experiments (> 200 cells were counted in each field and five fields were counted per experiment). (**G**) HEK293 cells stably expressing the cGAS delN with LL/RK mutation (delN-LL/RK) were stimulated with or without 1 μg/ml ctDNA for 4□h. Then, the cell lysates were fractionated into five fractions: cytoplasmic, membrane, nuclear soluble, chromatin-bound, and cytoskeletal. The fractions were blotted as indicated. C1QBP: membrane marker; TBK1 and α-tubilin: cytosolic marker; H3: nuclear marker.

Next, we examined endogenous cGAS localization in cells by immunofluorescence assays (IFA). We used a newly developed anti-human cGAS antibody by Cell Signaling Technology and validated it by western blotting (Fig. S1B) and IFA (Fig. 1C) in cGAS wild type and knockout H1299 cells. Using this antibody, we found that cGAS localized in the cytosol, nucleus, nucleoli, micronuclei, chromosome, chromatin bridge, and perinuclear region (Fig. 1D). Furthermore, we examined the endogenous cGAS localization in several other cell lines. However, the patterns of endogenous cGAS distribution varied in different cell lines, irrespective of cytosolic DNA stimulation (Figs. 1C, S1C-S1F). In agreement with previous studies, we found that endogenous cGAS localized in the nucleus in all tested cell lines, suggesting a potential role of nuclear cGAS.

### The N-terminus and NES regulate cGAS nuclear localization

Previous efforts have been made to determine the region responsible for cGAS nuclear localization. One study suggested that the region of amino acids 161-212 is essential for cytosolic retention of human cGAS (Gentili et al., 2019). There is a conserved classic nuclear export signal (NES) “**<u>L</u>**xxx**<u>L</u>**xx**<u>L</u>**x**<u>L/I</u>**” within amino acids 161-212 (Fig. S2A) (Sun et al., 2021). To examine the role of the NES in cGAS subcellular localization, we deleted the NES in cGAS. IFA assays showed that the NES deletion led to a slight increase of nuclear cGAS (Fig. S2B). We further mutated two leucines in the NES into arginine and lysine (LL/RK) in mouse cGAS, respectively. These mutations not only disrupt the NES but also converted the NES into a nuclear localization signal (NLS) (Fig. 1E). IFA assays showed that the LL/RK mutation dramatically increased cGAS nuclear localization (Fig. 1F). However, there were still approximately 37% of cells in which cGAS resided in both the nucleus and the cytoplasm (Fig. 1F). These data suggest that the NES might be required but not sufficient for cGAS cytosolic localization.

We further examined which domain might control cGAS nuclear localization (Fig. S2A). We stably transfected the N-terminal domain (N) and the N deletion mutant (delN) of cGAS into HEK293 cells. Subcellular fractionation found that the N and delN showed distinct localizations (Fig. S2C). The N was mainly distributed in the cytosol with a small portion in the nuclear soluble fraction. As reported recently (Li et al., 2021), the delN localized in the membrane fraction due to the exposure of mitochondrial targeting signal (MTS) (Fig. S2A). Furthermore, delN was also found in the chromatin-bound and cytoskeletal fractions (Fig. S2C). IFA assays further corroborated the cytosolic localization of the N and the mitochondrial and nuclear localizations of delN (Fig. S2D), suggesting that the N-terminus is also involved in cGAS cytosolic localization. Thus, we mutated the two leucines in the NES into arginine and lysine in delN of cGAS (delN-LL/RK) to disrupt the NES and MTS. Interestingly, subcellular fractionation found that the delN-LL/RK mutant is exclusively expressed in the chromatin-bound fraction (Fig. 1G). Like endogenous cGAS, cytosolic DNA stimulation had little effect on the localization of delN-LL/RK (Fig. 1G). Overall, our data suggest that both NES and the N-terminal domain regulate cGAS subcellular localization.

### HSV-1 infection causes cGAS release from the chromatin to the nuclear soluble fraction

As most DNA viruses are uncoated and then replicate in the nucleus, it would be the best opportunity for the host to detect viral DNA in the nucleus at the early stage of viral infection. However, nuclear cGAS is immobilized on the chromatin. We hypothesized that DNA virus infection in the nucleus might cause cGAS release from the chromatin. In this regard, we infected RAW 264.7 cells with herpes simplex virus type 1 (HSV-1), a DNA virus that replicates in the nucleus. As shown in Figure 2A, HSV-1 infection caused a portion of cGAS to translocate to the nuclear soluble fraction. By contrast, cytosolic DNA stimulation has little effect on cGAS subcellular localization in RAW 264.7 macrophages (Fig. S3A). Furthermore, we also examined three other viruses, vaccinia virus (VACV), influenza A viruses (IAV), and vesicular stomatitis virus (VSV). VACV is a DNA virus that replicates in the cytoplasm. IAV and VSV are RNA viruses and replicate in the nucleus and the cytosol, respectively. However, all these viruses failed to induce cGAS translocation to the nuclear soluble fraction in RAW 264.7 macrophages (Figs. S3B-S3D), suggesting that DNA virus replication in the nucleus is required for cGAS release from the chromatin to the nuclear soluble fraction.

**Figure. 2.**
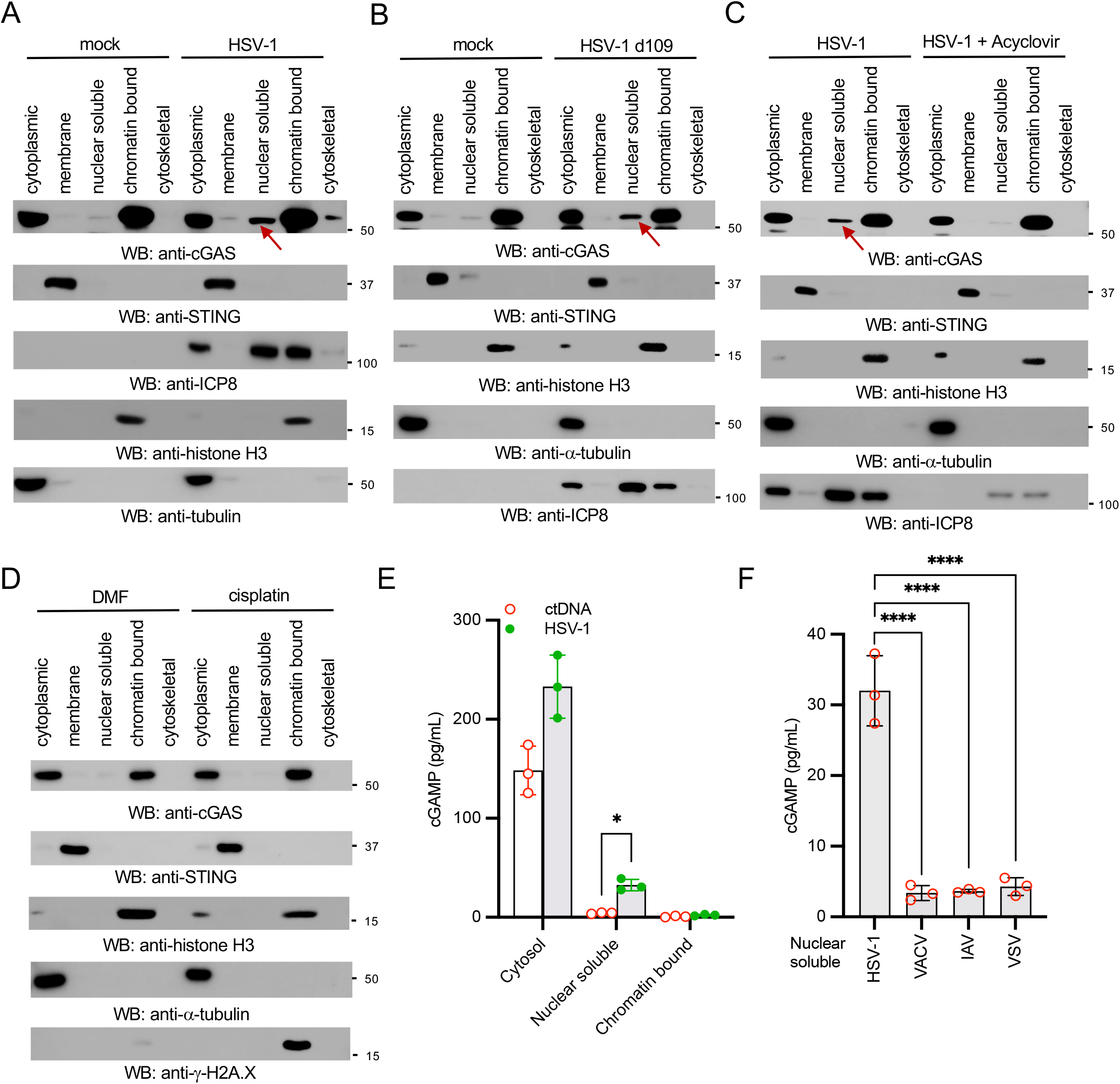
HSV-1 infection causes cGAS release from the chromatin to the nuclear soluble fraction. (**A**) RAW 264.7 macrophages were infected with HSV-1 for 16 h, and then the subcellular fractions of cell lysates were blotted as indicated. The arrow indicates nuclear soluble cGAS after viral infection. (**B**) RAW 264.7 cells were mock-infected or infected with 0.1 MOI of HSV-1 d109 for 16 h. Cell lysates were fractionated and blotted as indicated. The arrow indicates nuclear soluble cGAS after viral infection. (**C**) RAW 264.7 cells were pretreated without or with 8 μg/ml of acyclovir for 16 h, followed by infection with 0.1 MOI of HSV-1 for 16 h. Cell lysates were fractionated and blotted as indicated. The arrow indicates nuclear soluble cGAS. (**D**) RAW 264.7 macrophages were treated with dimethylformamide (DMF) or 50 μM cisplatin for 3 h. Cell lysates were fractionated and blotted as indicated. STING: membrane marker; α-tubulin: cytosolic marker; H3: nuclear marker; ICP8: HSV-1 viral protein; γ-H2A.X: DNA damage marker. (**E**) RAW 264.7 cells were transfected with ctDNA or infected with HSV-1 for 8 h. The cytoplasmic, nuclear soluble and chromatin-bound fractions were harvested for *in vitro* cGAS enzymatic activity assays. (**F**) RAW 264.7 cells were infected with HSV-1, VACV, IAV, or VSV for 8 h. The nuclear soluble extracts were harvested for *in vitro* cGAS enzymatic activity assays.

### Inhibiting HSV-1 replication blocks cGAS release from the chromatin

To determine the mechanism by which HSV-1 induces cGAS release from the chromatin, we first examined the effects of HSV-1 d109 mutant virus in which the IFN-suppression viral genes are deleted (Eidson et al., 2002). As shown in Figure 2B, the d109 mutant also induced nuclear soluble cGAS, suggesting these IFN-suppression viral proteins have little effect on cGAS localization. Next, we examined the effects of viral replication on cGAS release. The HSV-1 DNA polymerase inhibitor acyclovir was used to inhibit viral replication. As predicted, viral protein ICP8 expression reduced after acyclovir treatment. More interestingly, acyclovir blocked viral infection-induced nuclear soluble cGAS (Fig. 2C).

As HSV-1 infection can cause nuclear stress and host DNA damage, we suspected that DNA damage might induce the release of cGAS from chromatin tethering. In this regard, we treated RAW 264.7 macrophages with a DNA damage agent, cisplatin (Hu et al., 2016). As shown in Figure 2D, cisplatin induced□γH2AX (phosphorylated S139 H2AX histone), a hallmark of DNA damage. However, cisplatin failed to induce nuclear soluble cGAS (Fig. 2D). Taken together, these data suggest that DNA virus nuclear replication, but not IFN-suppression viral proteins or DNA damage, causes cGAS release from the chromatin.

### Nuclear soluble cGAS is constitutively active

It has been reported that R241E mutation results in the release of cGAS from the chromatin (Volkman, 2019). We stably expressed cGAS R241E mutant in HEK293 cells that are lacking endogenous cGAS. As reported previously (Volkman, 2019), the R241E mutant was only present in the cytosolic and the nuclear soluble fractions (Fig. S3E) and produced a significant amount of cGAMP without DNA ligand stimulation (Fig. S3F), suggesting the nuclear soluble cGAS is constitutively active.

Next, we examined whether the nuclear soluble cGAS induced by HSV-1 was active. We treated RAW 264.7 cells with calf thymus DNA (ctDNA) and HSV-1 followed by subcellular fractionation. As shown in Figure 2E, the cytosolic cGAS had comparable enzymatic activities in cells infected with HSV-1 and treated with ctDNA. However, in vitro cGAS enzymatic assays showed that a significantly higher level of cGAMP was produced by the nuclear soluble extract of the HSV-1-infected RAW 264.7 cells but not the ctDNA-treated cells (Fig. 2E), suggesting the active state of the nuclear soluble cGAS. Furthermore, we compared the cGAS activity in the nuclear soluble fraction in cells infected with different viruses. As shown in Figure 2F, cGAMP production was only detected in the nuclear soluble extract from cells infected with HSV-1, but not other viruses. These data are consistent with the subcellular fraction results that IAV, VSV, and VACV failed to induce nuclear soluble cGAS.

### Establish a cell line exclusively expressing nuclear cGAS

To obtain a cell line exclusively expressing nuclear cGAS, we fused the NLS of SV40 large T-to the C-terminus of cGAS (cGAS-NLS) and the LL/RK mutant (LL/RK-NLS). We then stably transfected these constructs into HEK293 cells. As shown in Figure S4A, although most cGAS-NLS proteins resided in the nuclear fraction, there was still a small amount of cGAS in the cytosol. However, the LL/RK-NLS was exclusively in the nuclear fractions (Fig. S4A). To further exclude newly synthesized cytosolic cGAS, we generated an inducible system for cGAS, cGAS-NLS, and the LL/RK-NLS (Fig. 3A) and stably transfected them into HEK293 cells (Fig. S4B). As shown in Figure S4C, cGAS proteins were stable within 8 h after doxycycline removal. Furthermore, the induced LL/RK-NLS protein exclusively resided in the nucleus by immunofluorescence assays (Fig. S4D).

**Figure. 3.**
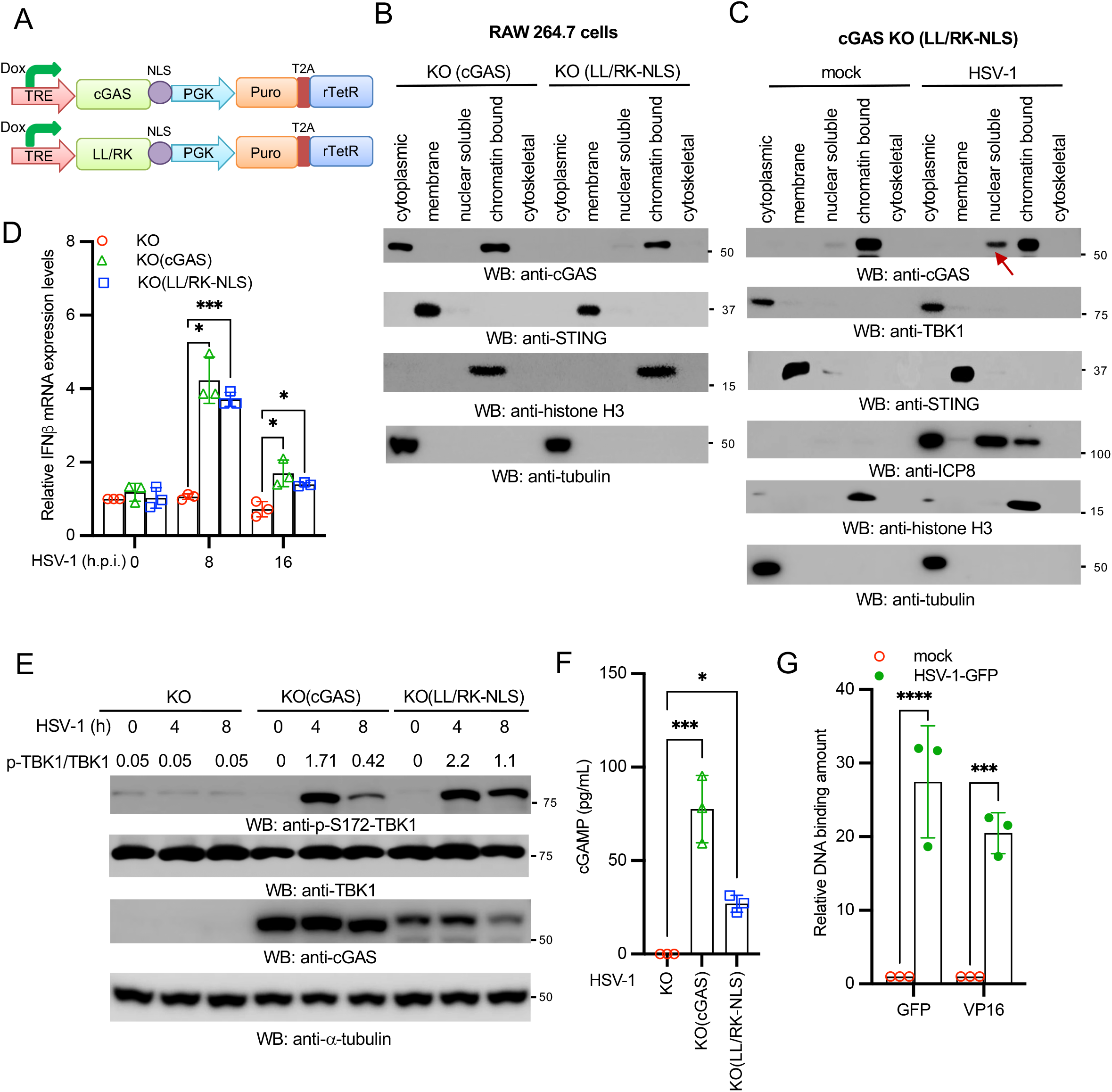
Nuclear soluble cGAS senses HSV-1 and instigates innate immune response. (**A**) Schematic of the doxycycline (Dox)-induced cGAS-NLS and LL/RK-NLS constructs. TRE: Tet Response Element; Puro: puromycin; T2A: Thosea asigna virus 2A-like peptide. (**B**) cGAS RAW 264.7 knockout macrophages reconstituted with the inducible cGAS or the inducible LL/RK-NLS mutant were fractionated into five fractions and blotted as indicated. STING: membrane marker; α-tubulin: cytosolic marker; ICP8: HSV-1 viral protein; H3: nuclear marker. (**C**) The cGAS KO (LL/RK-NLS) RAW 264.7 cells were mock-infected or infected with HSV-1 for 16 h. Then, cells were fractionated into five fractions and blotted as indicated. STING: membrane marker; TBK1 and α-tubulin: cytosolic marker; H3: nuclear marker. Red arrow indicates nuclear soluble cGAS. (**D**) The cGAS KO, KO (cGAS), and KO (LL/RK-NLS) RAW 264.7 cells were infected with 1 MOI of HSV-1 d109 for designated hours post-infection (h.p.i.). Real-time PCR was performed to determine the relative IFNβ mRNA levels. Data represent means ± s.d. of three independent experiments. All experiments were repeated three times. The *P*-value was calculated by two-way ANOVA followed by Dunnett’s multiple comparisons test (* *P* < 0.05, *** *P* < 0.001). (**E**) The cGAS KO, KO (cGAS), and KO (LL/RK-NLS) RAW 264.7 macrophages were infected with 1 MOI of HSV-1 for indicated times. Cell lysates were collected and blotted as indicated. Band densitometry was calculated by Image J. The relative ratio of phosphorylated TBK1 to total TBK1 in each lane was indicated. (**F**) cGAS KO, KO (cGAS), and KO (LL/RK-NLS) RAW 264.7 macrophages were infected with 1 MOI of HSV-1 for 8 h. The amount of cGAMP in each cell line was determined by ELISA assays. All experiments were repeated three times. The *P* value was calculated by one-way ANOVA followed by Tukey’s multiple comparisons test (* *P* < 0.05, *** *P* < 0.001). (**G**) HEK293 cells stably expressing FLAG tagged LL/RK-NLS mutant were infected with HSV-1-GFP for 8 h. ChIP assays were performed using the anti-FLAG antibody. Real-time PCR was performed to determine the relative binding amount of GFP and viral gene VP16. Data represent means ± s.d. of three independent experiments. All experiments were repeated three times. The *P*-value was calculated by two-way ANOVA followed by Dunnett’s multiple comparisons test (*** *P* < 0.001, **** *P* < 0.0001).

To determine the functional role of nuclear soluble cGAS, we applied the inducible cGAS expression system into RAW 264.7 knockout macrophages. We generated cGAS knockout in RAW 264.7 macrophages (Fig. S4E). cGAS knockout cells failed to produce IFNβ and ISGs, such as IP10 and RANTES, when infected with HSV-1 (Fig. S4F). Next, we reconstituted the inducible cGAS and the LL/RK-NLS mutant in cGAS knockout RAW 264.7 macrophages (hereinafter referred to as KO (cGAS) and KO (LL/RK-NLS), respectively) (Fig. S4G). Subcellular fractionation showed that LL/RK-NLS, but not the wild type cGAS, only resided in the chromatin-bound extract of the reconstituted cells after doxycycline induction (Fig. 3B).

### Nuclear soluble cGAS senses HSV-1 and activates innate immune responses

To examine the role of nuclear cGAS in HSV-1 infection, we infected the cGAS KO (LL/RK-NLS) RAW 264.7 cells with HSV-1. HSV-1 infection induced cGAS release from the chromatin to the nuclear soluble (Fig. 3C), which further corroborates that the nuclear soluble cGAS comes from the chromatin. Next, we compared innate immune responses to HSV-1 d019 in the cGAS KO, KO (cGAS), and KO (LL/RK-NLS) RAW 264.7 cells. As expected, cGAS KO cells failed to respond to HSV-1 infection; however, cGAS and LL/RK-NLS restored the mRNA expression of IFNβ, RANTES, and IP10 (Figs. 3D, S4H-S4I). The reconstitution of cGAS and LL/RK-NLS also rescued TBK1 phosphorylation, the hallmark of activation of IFN production pathways (Fig. 3E). Furthermore, KO (LL/RK-NLS) RAW 264.7 cells produced a significant amount of cGAMP after HSV-1 infection (Fig. 3F), suggesting that nuclear soluble cGAS is activated by HSV-1 infection.

It has been reported that cGAS can bind host genomic DNA (Gentili et al., 2019); however, whether nuclear soluble cGAS binds viral DNA is unknown. Thus, we infected the KO (LL/RK-NLS) cells with the HSV-1 carrying a GFP. After infection, the nuclear soluble extracts were subject to chromatin immunoprecipitation (ChIP) assay. ChIP assays found that cGAS bound GFP and HSV-1 VP16 DNA (Fig. 3G), suggesting that cGAS can sense HSV-1 DNA in the nuclear soluble fraction.

### Nuclear cGAS inhibits HSV-1 infection

We examined whether nuclear soluble cGAS inhibits HSV-1 infection. In this regard, cGAS KO, KO (cGAS), and KO (LL/RK-NLS) cells were infected with HSV-1-GFP. As shown in Figure 4A, wild type cGAS and the LL/RK-NLS mutant rescued the antiviral activity in cGAS knockout cells, evidenced by the reduced number of GFP staining cells. Consistently, the reconstitution of wild type cGAS or the LL/RK-NLS mutant also reduced the expression of viral RNA (Fig. 4B), viral protein (Fig. 4C), and the production of viral particles (Fig. 4D), suggesting that nuclear soluble cGAS limits HSV-1 infection. Furthermore, we examined the role of nuclear cGAS in other viruses. We infected cGAS KO, KO (cGAS), and KO (LL/RK-NLS) cells with HSV-1, VACV, VSV, and IAV reporter viruses. The infection activity of HSV-1, but not other viruses, reduced in the KO (LL/RK-NLS) cells (Fig. 4E), which is consistent with that only HSV-1 induces nuclear soluble cGAS. Interestingly, the reconstitution of wild type cGAS not only restricted HSV-1 infection but also impaired the infection of VACV and VSV (Fig. 4E). As VSV and VACV replicate in the cytoplasm, these data suggest that cytosolic cGAS is critical for host defense to cytosolic pathogens. Taken together, our data demonstrate that nuclear soluble cGAS can sense DNA virus infection in the nucleus, instigate innate immune response, and inhibit viral infection.

**Figure. 4.**
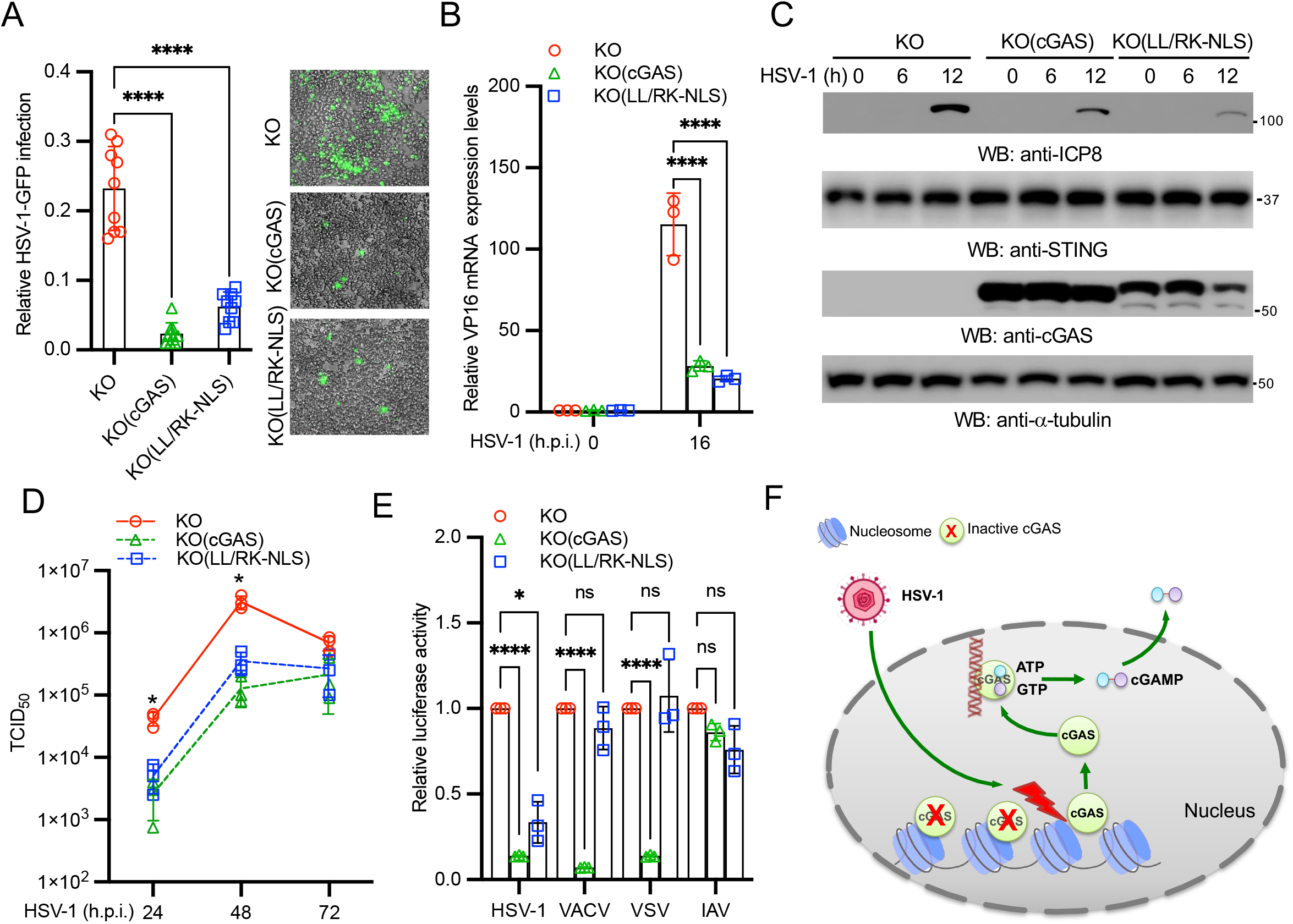
Nuclear soluble cGAS inhibits HSV-1 infection. (**A**) cGAS KO, KO (cGAS), KO (LL/RK-NLS) RAW 264.7 cells were infected with 0.1 MOI of HSV-1-GFP for 16□h. Cells expressing GFP were examined and counted under a fluorescence microscope. The relative infection was determined by the ratio of positive cells. Data represent means ± s.d. of three independent experiments (> 200 cells were counted in each field and five fields were counted per experiment). The *P*-value was calculated by one-way ANOVA followed by Dunnett’s multiple comparisons test (**** *P* < 0.0001). (**B**) The cGAS KO, KO (cGAS), KO (LL/RK-NLS) RAW 264.7 cells were infected with 1 MOI of HSV-1 for 16□h. Then, cells were collected for RNA extraction. The RNA levels of HSV-1 VP16 were determined by real-time PCR. All experiments were repeated three times. The *P*-value was calculated by two-way ANOVA followed by Dunnett’s multiple comparisons test (**** *P* < 0.0001). (**C**) The cGAS KO, KO (cGAS), KO (LL/RK-NLS) RAW 264.7 macrophages were infected with 1 MOI of HSV-1 for indicated times. Cell lysates were collected and blotted as indicated. ICP8 is an HSV-1 viral protein. (**D**) The cGAS KO, KO (cGAS), KO (LL/RK-NLS) RAW 264.7 macrophages were infected with 0.01 MOI of HSV-1 for the indicated days. HSV-1 titers were determined in Vero cells. Data represent means ± s.d. of three independent experiments. All experiments were repeated three times. The *P*-value was calculated by two-way ANOVA followed by Dunnett’s multiple comparisons test (* *P* < 0.05 vs. the cGAS KO cells). (**E**) The cGAS KO, KO (cGAS), KO (LL/RK-NLS) RAW 264.7 macrophages were infected with HSV-1-Luc, VACV-Luc, VSV-Luc, or IAV-Gluc for 16 h. Luciferase activities were measured to determine the relative infection activity. Data represent means ± s.d. of three independent experiments. All experiments were repeated three times. The *P*-value was calculated by two-way ANOVA followed by Dunnett’s multiple comparisons test (* *P* < 0.05, **** *P* < 0.0001, ns: not significant). (**F**) Model of nuclear soluble cGAS sensing DNA virus infection.

## DISCUSSION

It has been reported that cGAS locates predominantly in the cytoplasm (Sun et al., 2013), predominantly in the nucleus (Volkman, 2019), on the plasma membrane (Barnett et al., 2019), in both the cytoplasm and the nucleus (Gentili et al., 2019), mitosis-associated nuclear localization (Gentili et al., 2019; Yang et al., 2017), or phosphorylation-mediated nuclear import (Liu et al., 2018). Although the discrepancy is partially due to the cell type, the major reason is that most of the conclusions are based on imaging of the epitope-tagged to cGAS. Most commercial antibodies are not suitable for immunostaining and failed to pass the validation using cGAS knockout human and mouse cells in our hands. A recently developed human cGAS antibody by Cell Signaling Technology passed our in-house validation; therefore, we revisited the subcellular localization of endogenous cGAS by using the validated utility. Consistent with previous studies, we found endogenous cGAS localized in the cytoplasm, nucleus, chromosome, chromatin bridge, and micronuclei. However, different cell lines showed distinct subcellular localization patterns. For example, the localization pattern of cGAS is almost similar in most HFF-1 cells but cGAS could be either cytosolic or nuclear in H1299 cells. Our data imply additional mechanisms might be required to regulate endogenous cGAS localization in different cells.

Currently, it is not clear how cGAS enters the nucleus. One study proposed that cGAS nuclear localization results from nuclear envelope breakdown in mitosis or nuclear envelope rupture in interphase (Gentili et al., 2019). Another study reported that the export of nuclear cGAS to the cytosol is required for cytosolic DNA sensing based on the observation of accumulation of cytosolic cGAS after DNA stimulation (Sun et al., 2021). Although the mutation of the NES moderately altered cGAS subcellular localizations, we did not observe a significant accumulation of cytosolic cGAS in multiple cell lines after DNA stimulation. However, our approaches cannot exclude the nucleocytoplasmic shuttling of cGAS. Nonetheless, there is a fair amount of cytosolic cGAS present in the cytosol of unstimulated cells. Another recent study showed that the collided ribosomes induced nuclear cGAS translocation to the cytosol under translation stress (Wan et al., 2021). The critical question for these models is how cytosolic DNA or translation stress transduces a signal to nuclear cGAS proteins which are chromatin-bound. Logically, chromatin-bound cGAS would be first released into the nuclear soluble fraction, like the R222E and R241E mutants. Then, the nuclear soluble cGAS is exported into the cytosol. However, nuclear soluble cGAS is barely seen during cytosolic DNA stimulation. Whether and how cytosolic DNA stimulation induces cGAS nuclear export needs further investigation in the future.

It is intriguing to know why cGAS must localize in the nucleus, even it is a sensor that detects cytosolic DNA. One plausible explanation is that the nuclear cGAS senses the viruses or other invading pathogens that can evade the surveillance of cytosolic cGAS by uncoating in the nucleus. Indeed, most DNA viruses, like HSV-1, replicate in the nucleus and expose viral DNA in the nucleus during early infection. Paradoxically, to avoid being constitutively activated in the nucleus, cGAS is tethered to the chromatin. Our study now reveals that HSV-1 infection induces the release of cGAS from the chromatin to the nuclear soluble fraction. The “free” nuclear cGAS is in a DNA-rich environment, and the untethering is sufficient to activate cGAS by either host or viral DNA (Fig. 4F). Taken together, our study demonstrates that nuclear soluble cGAS is critical for sensing viral DNA in the nucleus at the early stage of DNA virus infection.

## MATERIALS AND METHODS

### Cells

HEK293 cells (ATCC, # CRL-1573), RAW 264.7 (ATCC, # TIB-71), NCl-H596 cells (ATCC, HTB-178), HFF-1 (ATCC, # SCRC-1041), MDA-MB-231 (Sigma, 92020424-1VL), and Vero cells (ATCC, # CCL-81) were maintained in Dulbecco’s Modified Eagle Medium (Life Technologies, # 11995-065) containing antibiotics (Life Technologies, # 15140-122) and 10% fetal bovine serum (Life Technologies, # 26140-079). NCI-H1299 cells (ATCC, # CRL-5083), THP-1 cells (ATCC) were cultured in RPMI Medium 1640 (Life Technologies, # 11875-093) plus 10% fetal bovine serum. A549 cells (ATCC, # CCL-185) were cultured in RPMI Medium 1640 (Life Technologies, # 11875-093) plus 10% fetal bovine serum and 1 × MEM Non-Essential Amino Acids Solution (Life Technologies, # 11140-050).

### Viruses

HSV-1 KOS strain was purchased from ATCC (#VR-1493). Viral titration was performed as the following. Vero cells were infected with a serial diluted HSV-1. After 1□h, the medium was removed and replaced by the DMEM plus 5% FBS and 1% agarose. Cells were examined for cytopathic effects to determine TCID50 or were fixed using the methanol–acetic acid (3:1) fixative and stained using a Coomassie blue solution to determine MOI.

### Plasmids

Mouse and human cGAS cDNA were synthesized and cloned into pCMV-3Tag-8 to generate mcGAS-FLAG and hcGAS-FLAG. Point mutations and deletions of cGAS were constructed using a Q5® Site-Directed Mutagenesis Kit (New England Biolabs, # E0554S).

### Antibodies

Primary antibodies: Anti-β-actin (Abcam, # ab8227), anti-FLAG (Sigma, # F3165), anti-TBK1 (Cell Signaling Technology, # 3504S], anti-phospho-TBK1 (Ser172) (Cell Signaling Technology, # 5483S), anti-C1QBP (Cell Signaling Technology, # 6502S), anti-α-Tubulin (Cell Signaling Technology, # 3873S), anti-Histone H3 (Cell Signaling Technology, # 4499S), anti-mouse STING (Cell Signaling Technology, # 50494S), anti-human STING (R&D Systems, # MAB7169-SP), anti-vimentin (R&D Systems, # MAB21052-SP), anti-ICP8 (Abcam, # ab20194), anti-NP (GenScript, # A01506-40), anti-γ-H2AX (ABclonal, # AP0099), anti-human cGAS (Cell Signaling Technology, #79978), anti-mouse cGAS (Cell Signaling Technology, #31659).

Secondary antibodies: Goat anti-Mouse IgG-HRP [Bethyl Laboratories, # A90-116P, WB (1:10,000)], Goat anti-Rabbit IgG-HRP [Bethyl Laboratories, # A120-201P, WB (1:10,000)], Alexa Fluor 594 Goat Anti-Mouse IgG (H+L) [Life Technologies, # A11005, IFA (1:200)], Alexa Fluor 488 Goat Anti-Rabbit IgG (H+L) [Life Technologies, # A11034, IFA (1:200)].

### Subcellular fractionation

Approximately 2 × 10^6^ cells were fractionated using the Subcellular Protein Fractionation Kit for Cultured Cells (Thermo scientific, #78840).

### CHIP-qPCR assay

Briefly, HEK293 cells stably expressing FLAG-tagged LL/RK-NLS were infected with HSV-1-GFP virus for 16 h. Cells were cross-linked with formaldehyde for 10 min, followed by adding 125 mM glycine to stop the cross-linking. Cells were then washed with 1 ml 1 x PBS two times and centrifuged for 5 min at 4°C, 1,000 x g. The pelleted cells were used for subcellular fractionation. The nuclear soluble extracts were immunoprecipitated with the anti-FLAG beads. After three times washings with 1 x PBS, cGAS complexes were eluted using FLAG peptides, and then the elutes were subject to qPCR assay.

### cGAMP assay

Cells were collected by centrifuging at 500 x g for 5 minutes, and then resuspended and lysed in PBS or the Immunoassay Buffer C (Cayman chemical, # 401703) through boiling at 95°C for 10 minutes. The lysates were then centrifuged for 30 min at 4 °C, 10,000 x g. Supernatants were collected for cGAMP ELISA assays. For subcellular fractions, the samples were diluted 10 x with the Immunoassay Buffer C (Cayman chemical, # 401703). The cGAMP amount was determined by ELISA assays according to the manufacture’s protocols (Cayman chemical, # 501700).

### Sample preparation, Western blotting, and immunoprecipitation

Approximately 1 × 10^6^ cells were lysed in 500 µl of tandem affinity purification (TAP) lysis buffer [50 mM Tris-HCl (pH 7.5), 10 mM MgCl_2_, 100 mM NaCl, 0.5% Nonidet P40, 10% glycerol, the Complete EDTA-free protease inhibitor cocktail tablets (Roche, # 11873580001)] for 30 min at 4 □°C. The lysates were then centrifuged for 30 min at 15,000 rpm. Supernatants were mixed with the Lane Marker Reducing Sample Buffer (Thermo Fisher Scientific, # 39000) and boiled at 95 °C for 5 minutes.

Western blotting and immunoprecipitation were performed as described in a previous study (Zhao et al., 2019). Briefly, samples (10–15□μl) were loaded into Mini-Protean TGX Precast Gels, 15 well (Bio-Rad, # 456-103), and run in 1 × Tris/Glycine/SDS Buffer (Bio-Rad, # 161-0732) for 60□min at 140□V. Protein samples were transferred to Immun-Blot PVDF Membranes (Bio-Rad, # 162-0177) in 1 × Tris/Glycine buffer (Bio-Rad, # 161-0734) at 70□V for 60□min. PVDF membranes were blocked in 1 × TBS buffer (Bio-Rad, # 170-6435) containing 5% Blotting-Grade Blocker (Bio-Rad, # 170-6404) for 1□h. After washing with 1 × TBS buffer for a total of 30□min (10 min each time, repeat 3 times), the membrane blot was incubated with the appropriately diluted primary antibody in antibody dilution buffer (1 × TBS, 5% BSA, 0.02% sodium azide) at 4□°C for 16□h. Then, the blot was washed three times with 1 × TBS (each time for 10□min) and incubated with secondary HRP-conjugated antibody in antibody dilution buffer (1:10,000 dilution) at room temperature for 1□h. After three washes with 1 × TBS (each time for 10□min), the blot was incubated with Clarity Western ECL Substrate (Bio-Rad, # 170-5060) for 1-2□min. The membrane was removed from the substrates and then exposed to the Amersham imager 600 (GE Healthcare Life Sciences, Marlborough, MA).

### Immunofluorescence assay

Cells were cultured in the Lab-Tek II CC2 Chamber Slide System 4-well (Thermo Fisher Scientific, # 154917). After the indicated treatment, the cells were fixed and permeabilized in cold methanol for 10□min at -20□°C. Then, the slides were washed with 1 × PBS for 10□min and blocked with Odyssey Blocking Buffer (LI-COR Biosciences, # 927-40000) for 1□h. The slides were incubated in Odyssey Blocking Buffer with appropriately diluted primary antibodies at 4□°C for 16□h. After 3 washes (10□min per wash) with 1 × PBS, the cells were incubated with the corresponding Alexa Fluor conjugated secondary antibodies (Life Technologies) for 1□h at room temperature. The slides were washed three times (10□min each time) with 1 × PBS and counterstained with 300□nM DAPI for 1□min, followed by washing with 1 × PBS for 1□min. After air-drying, the slides were sealed with Gold Seal Cover Glass (Electron Microscopy Sciences, # 3223) using Fluoro-gel (Electron Microscopy Sciences, # 17985-10).

### Real-time PCR

Total RNA was prepared using the RNeasy Mini Kit (Qiagen, # 74106). Five hundred nanograms of RNA was reverse transcribed into cDNA using the QuantiTect reverse transcription kit (Qiagen, # 205311). For one real-time reaction, 10 µl of SYBR Green PCR reaction mix (Eurogentec), including 100 ng of the synthesized cDNA plus an appropriate oligonucleotide primer pair, were analyzed on a 7500 Fast Real-time PCR System (Applied Biosystems). The comparative C*t* method was used to determine the relative mRNA expression of genes normalized by the housekeeping gene *GAPDH*. The primer sequences: mouse *Gapdh*, forward primer 5’-GCGGCACGTCAGATCCA -3’, reverse primer 5’-CATGGCCTTCCGTGTTCCTA -3’; mouse *Ifnb1*, forward primer 5’-CAGCTCCAAGAAAGGACGAAC -3’, reverse primer 5’-GGCAGTGTAACTCTTCTGCAT -3’; mouse *Cxcl10 (IP10)*, forward primer 5’-CCAAGTGCTGCCGTCATTTTC -3’, reverse primer 5’-GGCTCGCAGGGATGATTTCAA -3’; mouse *Ccl5 (RANTES)*, forward primer 5’-GCTGCTTTGCCTACCTCTCC -3’, reverse primer 5’-TCGAGTGACAAACACGACTGC-3’; GFP, forward primer 5’-AAGGGCATCGACTTCAAGG -3’, reverse primer 5’-TGCTTGTCGGCCATGATATAG -3’; HSV-1 VP16, forward primer 5’-GGACTGTATTCCAGCTTCAC -3’, reverse primer 5’-CGTCCTCGCCGTCTAAGTG -3’.

### Plasmid transfection

HEK293 and RAW cells were transfected using Lipofectamine 3000 or Lipofectamine LTX Transfection Reagent (Life Technologies, # L3000015) according to the manufacturer’s protocol.

### CRISPR/Cas9

The single guide RNA (sgRNA) targeting sequences: mouse *cGAS* sgRNA1: forward primer 5’-CACCGACGCAAAGATATCTCGGAGG -3’, reverse primer 5’-AAAC CCTCCGAGATATCTTTGCGTC-3’; sgRNA2 forward primer 5’-CACCG AGATCCGCGTAGAAGGACGA -3’; reverse primer 5’-AADCTCGTCCTTCTACGCGGATCTC -3’. SgRNA3 forward primer 5’-CACCGGCGGACGGCTTCTTAGCGCG -3’; reverse primer 5’-AAACCGCGCTAAGAAGCCGTCCGCC -3’. The sgRNA was cloned into lentiCRISPR v2 vector (Sanjana et al., 2014) (Addgene). The lentiviral construct was transfected with psPAX2 and pMD2G into HEK293T cells using PEI. After 72 h, the media containing lentivirus were collected. The targeted cells were infected with the media containing the lentivirus supplemented with 10 μg/ml polybrene. Cells were selected with 10 μg/ml puromycin for 14 days. Single clones were expanded for knockout confirmation by Western blotting.

### Stable Cell Line Selection

HEK293 cells and RAW 264.7 cells were transfected with the relative constructs or infected with the media containing the lentivirus supplemented with 10 μg/ml polybrene. Cells were selected with 200 μg/ml hygromycin or 10 μg/ml puromycin for 14 days. Stable cell lines were validated by Western blotting.

### Statistics and Reproducibility

The sample size was sufficient for data analyses. Data were statistically analyzed using the software GraphPad Prism 9. Significant differences between the indicated pairs of conditions are shown by asterisks (* *P* <0.05; ** *P* <0.01; *** *P* <0.001; **** *P* <0.0001).

## Author Contributions

SL conceived and supervised the project. SL, YW, KS, WH, and LW designed the study. YW, KS, WH, and LW performed the experiments. SL, YW, KS, WH, and LW analyzed the data. All authors contributed to manuscript writing, revision, read, and approved the submitted version.

## ACKNOWLEDGEMENT

This research was funded by the National Institutes of Health (R21AI137750 and R01AI141399 to S.L.).

## Competing interests

The authors declare no competing interests.

## Notes

### Competing Interest Statement

The authors have declared no competing interest.

